# Estimation of Brain Age Delta from Brain Imaging

**DOI:** 10.1101/560151

**Authors:** Stephen M. Smith, Diego Vidaurre, Fidel Alfaro-Almagro, Thomas E. Nichols, Karla L. Miller

## Abstract

It is of increasing interest to study “brain age” - the apparent age of a subject, as inferred from brain imaging data. The difference between brain age and actual age (the “delta”) is typically computed, reflecting deviation from the population norm. This therefore may reflect accelerated aging (positive delta) or resilience (negative delta) and has been found to be a useful correlate with factors such as disease and cognitive decline. However, although there has been a range of methods proposed for estimating brain age, there has been little study of the optimal ways of computing the delta. In this technical note we describe problems with the most common current approach, and present potential improvements. We evaluate different estimation methods on simulated and real data. We also find the strongest correlations of corrected brain age delta with 5,792 non-imaging variables (non-brain physical measures, life-factor measures, cognitive test scores, etc.), and also with 2,641 multimodal brain imaging-derived phenotypes, with data from 19,000 participants in UK Biobank.

## 1 Introduction

Brain imaging (and other sources of relevant data) can be used to predict “brain age” - the apparent age of individuals, when comparing their data against a population dataset spanning a range of ages. The difference between brain age and actual age (the “delta”) is often then computed, providing a measure of whether a subject’s brain appears to have aged more or less than the population average for their actual chronological age. For example, looking at structural MRI data, a high degree of atrophy (e.g., caused by disease) would cause a subject’s brain to appear older than a normal age-matched brain [Franke et al., 2010, Cole et al., 2017, Cole and Franke, 2017]

The approach typically taken is to use one or more imaging modalities, for example, acquiring a T1-weighted structural image from each subject. The data then receives some level of preprocessing, e.g., alignment to standard space and tissue type segmentation. The imaging data then becomes “features” for predicting brain age - for example, from voxelwise maps of grey matter partial volume estimates, the voxelwise values themselves can be the features. Alternatively, a smaller number of more highly-condensed features, such as volumes of grey matter within multiple distinct brain regions of interest, may be generated. The entire dataset of multiple subjects’ features, and their true ages, are fed into a supervised-learning algorithm (e.g., regression, support vector machine, deep learning), which learns to predict the subjects’ ages from their brain imaging features. The hope is that a given subject’s predicted brain age will deviate from their true age according to a meaningful delta, as long as this training is not badly overfitting.

While there have been a range of methods proposed for estimating brain age (i.e., choice of imaging-derived features to use, and choice of supervised-learning approach), there has been very little study of the optimal ways of then computing the delta. Presumably this is in part because this seems like a very simple calculation (delta equals brain age minus age). However, there are frequently various sources of bias in estimating brain age delta, which can give rise to significant false positives and false negatives when looking for associations between delta and other measures [Le et al., 2018]. Here we describe some important problems with this most common approach, and present potential improvements via explicitly laid out (albeit simple) mathematical frameworks.

We study these effects in simulated and real data, and suggest simple models for removing bias and increasing the accuracy of delta estimation. This includes models for correcting underestimation of brain aging, removing the resulting dependency of delta on age, modelling nonlinear dependence of brain aging (as a function of age), and studying non-additive brain age delta. One of the causes of bias (regression dilution, see below) has recently also been discussed in depth in [Le et al., 2018], where the same linear correction as Eq.4 below was proposed. A closely-related linear correction was also recently proposed in [Liang et al., 2019], although these authors suggest that regression towards the mean is the only source of bias, and we find empirically that non-Gaussian distribution of subject ages can be a major cause of bias.

## 2 Methods and Results

In order to present the clearest description of the gradual development of models throughout the paper, we do not separate out Methods and Results sections, but intermix discussions of model improvements with simulation and real-data results.

### 2.1 Common approach

We start with some basic definitions, and outline the typical approach for brain age delta estimation. Actual age is *Y* (an *N*_*subjects*_ ×1 vector), brain age is *Y*_*B*_ and the delta is *δ* = *Y*_*B*_ −*Y*. The imaging data matrix is *X*, which has *N*_*subjects*_ rows and *D* columns; the columns are features from the imaging data, and might be different voxels, or different IDPs (imaging-derived phenotypes - summary measures of brain structure and function).^1^

It is common to try to predict *Y*_*B*_ from *X*:

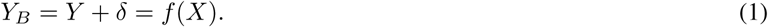

Although many alternatives have been proposed for the form of *f* () and the fitting of its model parameters, the issues explored in this paper generally hold, irrespective of these choices (e.g., most brain age literature shows underestimation of brain age for old subjects, and overestimation in young subjects). We start with a very simple linear model, with *f* (*X*) = *Xβ*, a multiple regression with parameters *β* (a *D* × 1 vector). We therefore have:

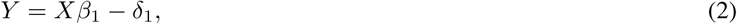

where we have added subscripts to differentiate some variables in this model from later variants. Clearly *δ*_1_ is the brain age delta being estimated, and the noise term in this formulation.

This multiple regression can be solved with standard simple methods,^2^for example, setting *β*_1_ = *X*^+^*Y* using the pseudo-inverse *X*^+^ = (*X*^*′*^*X*)^-1^*X*^*′*^. This gives the initially predicted brain age *Y*_*B*1_ = *XX*^+^*Y*, and brain age delta:

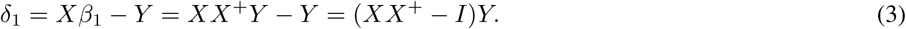

Two complications are immediately clear: *δ*_1_ will be orthogonal to the imaging data matrix *X* (though there is no reason to assume that this is desirable), and it will not be orthogonal to age (*Y*). This latter issue is likely to be problematic, given the simplest conception of the delta as being the difference between brain age and age, as a useful (objective, non-changing) description of a subject’s brain-health that is not dependent on their current age. In the extreme case of *X* being an entirely useless model for brain aging, *Xβ*_1_ *≈* 0 and *δ*_1_ will just be −1 times the actual age.

In addition, there are several factors that almost always cause the estimated *β*_1_ to be biased towards zero (Fig. 1), resulting in under-fitting to *Y* and hence “moving” age dependence into *δ*_1_:

**Figure 1:**
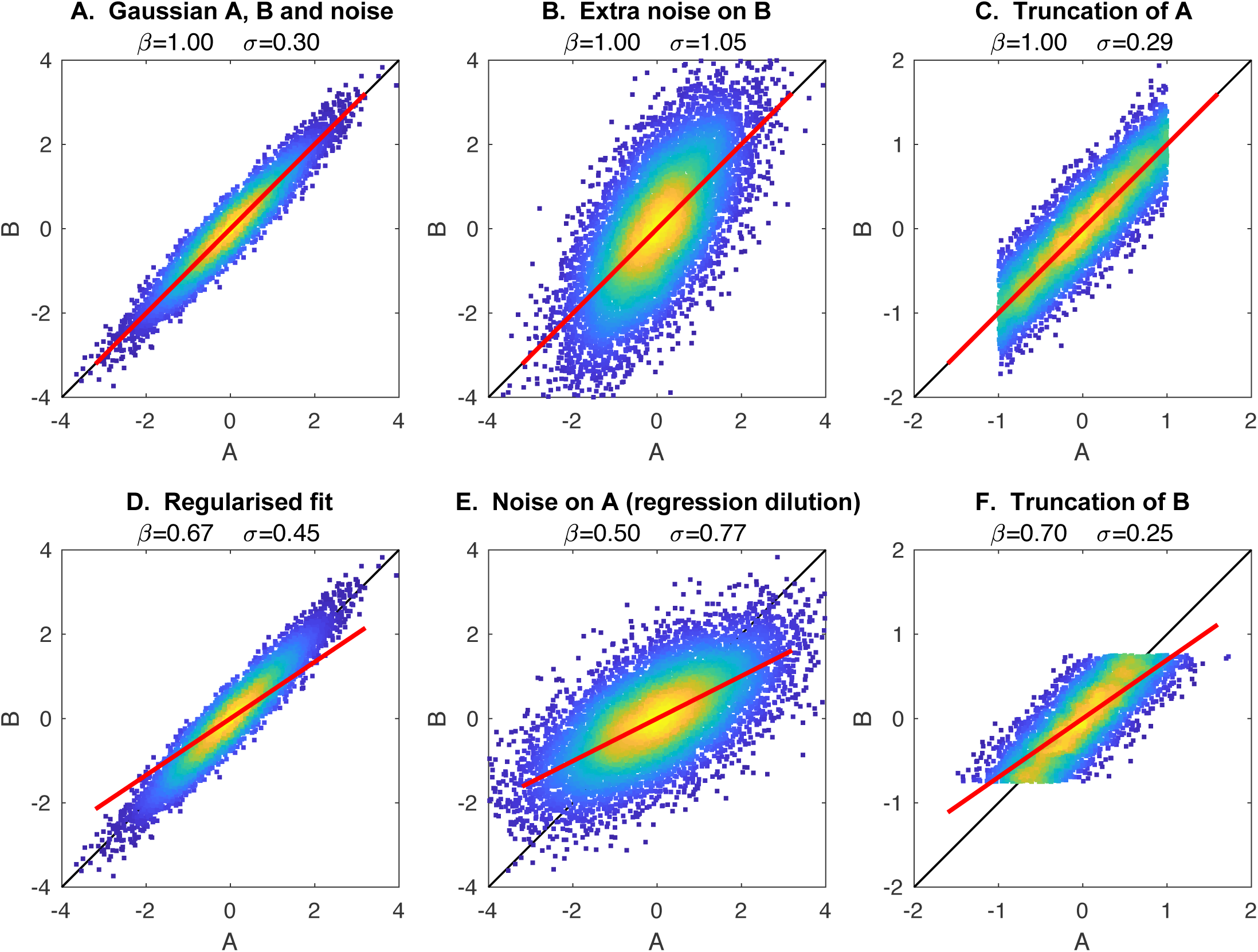
Examples of different causes of biased estimation of the model fit. The black line shows the correct model fit (*B* = *A*), while the red line shows the fitted model (*B* = *Aβ*). **A**. The model *A* is Gaussian distributed, and simulated *B* is set to be equal to *A*, with Gaussian noise added to *B*. The true model fit (*β* = 1) is correctly estimated. **B**. Increased measurement noise on *B* increases the error in the model fit (*σ*), but does not cause bias (even though the data cloud appears rotated). **C**. Changing *A* to being non-Gaussian does not harm the model fitting. **D**. Applying regularised model-fitting (here Tikhonov regularisation) biases the model fit towards zero (even though the data cloud is not rotated). **E**. Adding measurement noise to *A* (i.e., after it has been used to generate *B*) causes regression dilution. **F**. Truncating *B* (a special case of non-Gaussianity in *B*) causes a biased model fit.

1. It is typical for more sophisticated fitting methods (e.g., sparse/regularised regression) to result in underestimation of *β*_1_ [Grosenick et al., 2013].
2. It is common for datasets to not have a Gaussian distribution (across all subjects) for age, often because of hard limits on age in the study design. In the linear model framework, it does not generally matter what the distribution of the predictors is (here, *X*), but a non-Gaussian distribution in the independent variable (here *Y*) can cause serious bias (typically underestimation) in the estimated *β*_1_.
3. Errors in measurement of the predictors (*X*) also likely cause underestimation of *β*_1_ (“regression dilution”) [Le et al., 2018].

Note that the above formulations, as they stand, model the entire dataset together, to estimate brain age and the delta, and these causes of bias often exist in such a scenario. In practice, researchers often use cross-validation (or a group of healthy subjects and a clinically distinct subject group), where model parameters are learned from training data and then applied to left-out data in order to estimate delta, in a way that is more robust against model over-fitting. Applying such cross-validation (as opposed to all-in-one fitting) can be yet another factor causing non-orthogonality between estimated delta and age (because, like with regularisation, the tendency would be to increase under-fitting).

Finally, with *β*_1_ being underestimated, and age-dependence therefore moved into *δ*_1_, there is the danger that association tests against non-imaging variables (e.g., cognitive status, health outcomes) will be dominated by true age (i.e., be driven by the aging process), rather than the intended brain age delta. Obviously this can be eliminated through careful deconfounding of the non-imaging variables, but with age-dependence left in *δ*, this still results in loss of statistical sensitivity. To re-iterate the danger here: if (as seems to happen frequently in the literature) estimated brain age delta is not orthogonal to age, and if other variables (cognitive status, health measures, etc.) have not been deconfounded with respect to age, then any apparent associations between “brain age delta” and the non-imaging measures might be more driven by age and not true delta.

### 2.2 Stage-2 correction of delta

One very simple approach to correct for all of the above issues is to remove age dependence in *δ*_1_ in a second step. For now we will assume only linear corrections, and will return to nonlinear correction later.

The method is easily motivated by considering the scatterplot of *δ*_1_ against *Y* (Fig 2). As discussed above, ideally we would want the data to be distributed around *δ*_1_ = 0, with no overall slope (dependence on *Y*). If an overall slope is present, we can simply fit a straight line to the full data cloud and subtract this:

**Figure 2:**
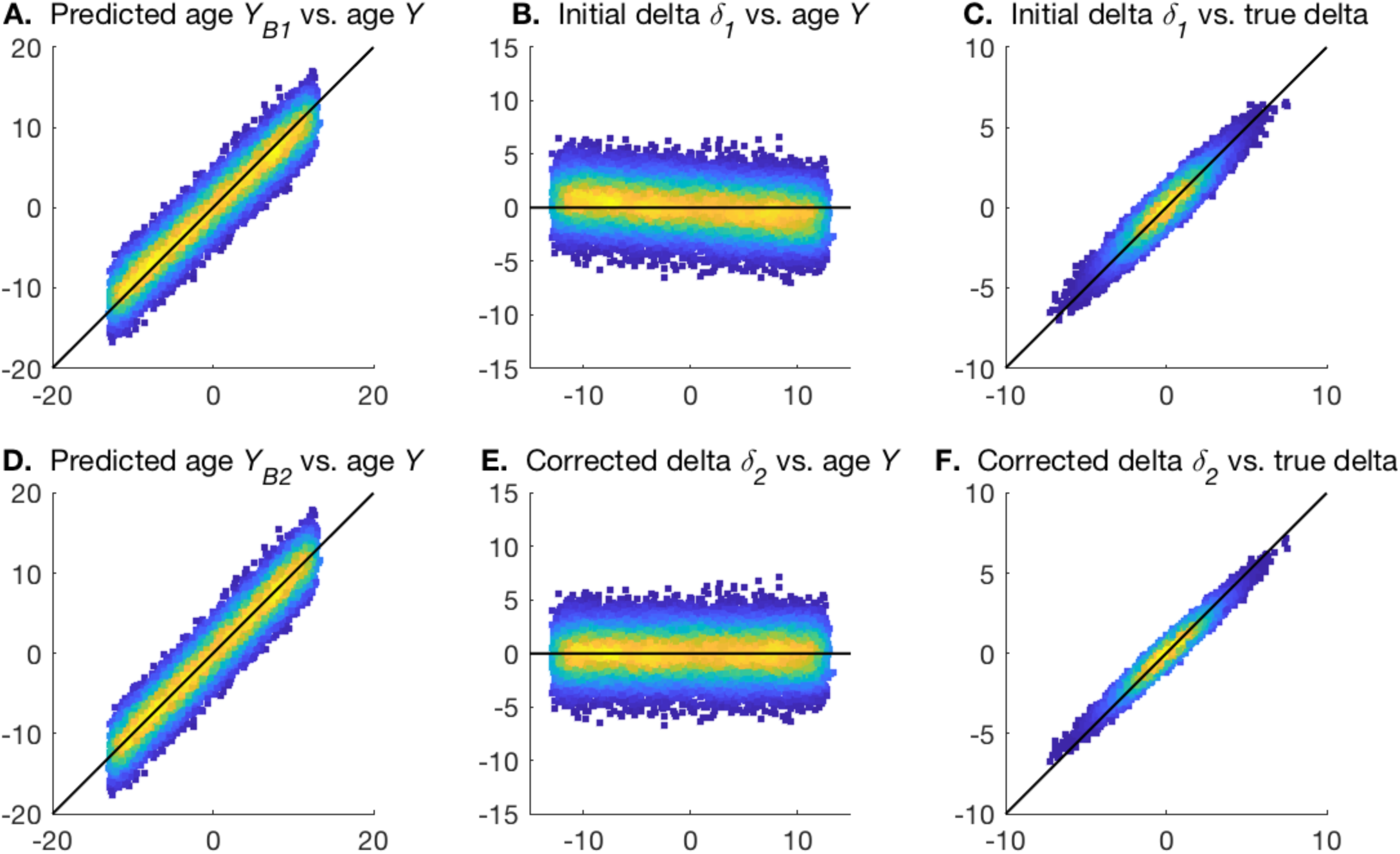
Examples of the two stages of age prediction. The black lines show the ideal unbiased model fits. **A,D**. The plots of predicted brain age from steps 1 and 2, vs. actual age. **B,E**. The plots of estimated brain age delta from steps 1 and 2, vs. actual age. **C,F**. The plots of estimated brain age delta from steps 1 and 2, vs. true delta.

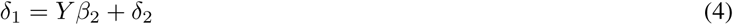

where we have defined *δ*_2_ (the residuals from this fitting), to be orthogonal to age, with biases (and modelling failures due to poor *X*) removed.

Although it is perfectly convenient (and easy to understand) to carry out this modelling in two separate steps, they can easily be combined into a single calculation:

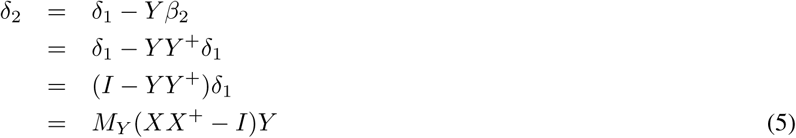

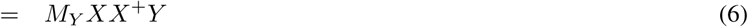

where *M*_*Y*_ = *I* −*Y Y* ^+^ is the “residual-forming matrix”, which orthogonalises a vector with respect to *Y*.^3^ The second term in the brackets in Eqn. 5 disappears because *M*_*Y*_ *Y* = 0. A simple intuitive interpretation of this is that *XX*^+^*Y* is the predicted age from step 1, and therefore *δ*_2_ is just the orthogonalisation of this with respect to actual age. The predicted brain age from this second step is *Y*_*B*2_ = *Y* + *δ*_2_.

Fig. 2 uses a simple simulation to illustrate the two stages of the model fitting described above. In **A** and **B** the biases in the one-stage modelling are apparent, and these are removed in **D** and **E**. Furthermore, the correlation of estimated delta with the true delta is improved in the second step.

### 2.3 Switching predictor matrix and age

Alternatively, one could frame the modelling in an arguably more natural framework, with *X* being dependent on *Y*_*B*_:

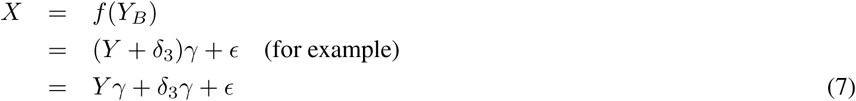

where *γ* is a 1 × *D* row vector, meaning that *Y γ* is a rank-1 approximation to *X*. This approach has the advantage of looking “causally sensible” (brain aging affects what we measure of the brain). Also, if this is solved using *Y* as the predictor variable, *δ*_3_ will be treated as being in the residuals and hence is defined as being orthogonal to age (and not to *X*). Finally, some problems such as regression dilution go away, as we can generally assume that there are no errors in *Y*.

However, one might (in general correctly) expect that this model is not as statistically powerful as the above approaches, given that each column in X is being modelled separately from each other (with respect to estimating *γ*).

Again, this formulation can be easily solved:

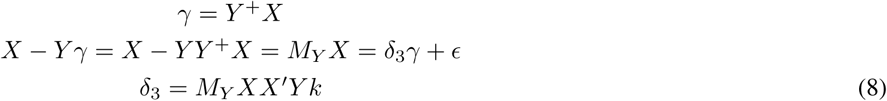

where scaling factor *k* = (*Y* ′*Y*)*/*(*Y* ′*XX*′*Y*), as derived from multiplying by the right-pseudo-inverse of *γ*. This defines the residuals *ϵ* as being vertically orthogonal to age, and horizontally orthogonal to *γ*.

It can immediately be seen that *δ*_3_ becomes equal to *kδ*_2_ if *X* is first orthonormalised before being used here (i.e., *X*^+^ = *X*′). This makes intuitive sense as we would now be saying that the columns in *X* do not depend on each other, so the potential for statistical insensitivity in the formulation of Eq.7 disappears.

### 2.4 Improved prediction accuracy via singular value decomposition

It is straightforward (and typical) for the first stage of the common approach to use regularised modelling applied to *X*, particularly given that *X* is frequently formed from (unwrapped) maps of voxels, resulting in there being too many variables in *X* for trivial application of multiple regression. Even if this were not the case, regularised estimations are known to often perform better than non-regularised estimations in terms of statistical efficiency, for a sufficiently good choice of the regularisation parameter.

As one option for regularisation, *X* can be pre-reduced using SVD (singular value decomposition): *X* = *USV* ^′^, where the *J* strongest eigen-subject-vectors from *U* (those that explain the strongest variance in *X*) would be used in place of *X*. This procedure is sometimes referred to as principal-component-regression [Massy, 1965, Franke et al., 2010], and has the general effect of “denoising” *X* and therefore improving overall modelling for an appropriate choice of *J*. By definition, *U* is orthonormal, and hence, as discussed above, this causes *δ*_2_ and *δ*_3_ to be the same (apart from the overall scaling). It is important to note that, even when using effective regularisation, it remains the case (as demonstrated in examples below) that the initial estimate (*δ*_1_) most commonly used in brain age studies remains suboptimal and biased. In the following sections we illustrate this with simulated and real data.

### 2.5 Cross-validation

Although not a primary focus of this paper, we briefly discuss here the use of cross-validation. In general, when there is any risk of an analysis over-fitting the data (e.g., where there is some data-dependence in the processing, such as is the case with any supervised machine learning), it is important to use methods such as cross-validation to avoid inflated estimates of modelling success.

The models covered in this paper are sufficiently well-conditioned when run on large subject numbers that cross-validation is not expected to change results greatly. In practice, we found that results reported in the various simulations reported below were only significantly altered (when using cross-validation) where the models fitted had the greatest numbers of features - i.e., when using the full (no SVD) model for *X*, or when using the maximum number of SVD eigenvectors. Nevertheless, all results reported below were estimated using cross-validation, which we now describe briefly.

We applied 10-fold cross-validation, where the data samples are randomly assigned into 10 roughly equal-sized groups. For each group of left out data, the other 90% of samples (subjects) are used to “train”, i.e., estimate model parameters. These parameters are then applied to the left-out subjects. In this case, the training refers to 4 analysis stages: a) confound removal (for real data), b) SVD reduction, c) *δ*_1_ initial estimation and d) *δ*_2*/*3_ correction.

Finally, note that if automated model tuning is to be carried out (e.g., optimal SVD dimensionality to be estimated from the data), then clearly this also should be done within a cross-validation framework.

### 2.6 Evaluations with simulations and real data

We now present results from simple but realistic simulations and real data.

#### Simulation 1

For Simulation 1, we set the number of samples (subjects) to 10000, set a fairly sharply truncated (non-Gaussian) distribution for the age range (total range approximately 45-75y), and added Gaussian *δ* with standard deviation 2y to form the gold-standard brain age. We then defined 100 underlying components (processes) of subject variation in “brain imaging” measures, the first being brain age, and the other 99 being random. We then mixed these ground truth population modes by a (100×3000) sparse mixing matrix (random Gaussian noise to the fifth power) to form 3000 imaging variables, resulting in an *X* of size 20000×3000. Finally, we standardised all columns in X to 1, and added measurement noise with a standard deviation of 0.5. (Reducing this noise to the kinder level of 0.1 does not make a large qualitative difference to the results.) We ran the simulation 20 times, and show the mean and standard deviation results across the 20.

The results can be seen in Table 1. The most important results are the 3 right-most columns, where (in the simulated datasets) *Q* shows how well the different estimates of *δ* correlate with the true *δ*. The first row shows results when the “raw” *X* is used (i.e., without SVD). This is outperformed by the best SVD-based analyses. However, even in this case it is clear that *δ*_1_ is not as accurate at recovering the true *δ* as *δ*_2_.

**Table 1:** Results from quantitative evaluations of brain age estimation. We show results for two different simulations and one real dataset. Different rows are for different SVD dimensionality reductions, with the first in each experiment being with no use of SVD at all. Columns 2-5 show the correlations of various estimated measures with true age. The next three show the mean absolute value of three different brain age *δ* estimates (for the true *δ* in the simulations, it is 1.6y). The final three columns (“*Q*”) show the most important aspect of the results: in the case of the simulations, these are the correlation between the true delta and the different estimations of delta. In the case of the real data from UK Biobank, where we do not know the true delta, we are using as a surrogate a summary measure of the significance of the correlation between estimated delta and 5792 non-imaging variables.

| SVD $J$ | $r(Y, \delta_1)$ | $r(Y, Y_{B1})$ | $r(Y, Y_{B2})$ | $r(Y, Y_{B3})$ | $\vert \delta_1\vert $ | $\vert \delta_2\vert $ | $\vert \delta_3\vert $ | $Q(\delta_1)$ | $Q(\delta_2)$ | $Q(\delta_3)$ |
| --- | --- | --- | --- | --- | --- | --- | --- | --- | --- | --- |
| Sim1 - | -0.23±0.00 | 0.94±0.00 | 0.95±0.00 | 0.94±0.00 | 2.0±0.0 | 1.9±0.0 | 2.0±0.1 | 0.74±0.01 | 0.76±0.01 | 0.80±0.03 |
| 1 | -0.99±0.01 | 0.06±0.10 | 0.99±0.01 | 0.06±0.09 | 6.2±0.1 | 0.4±0.5 | 537.1±645.4 | 0.00±0.01 | 0.02±0.03 | 0.02±0.03 |
| 10 | -0.94±0.08 | 0.29±0.18 | 0.96±0.03 | 0.29±0.18 | 5.8±0.7 | 1.5±0.7 | 22.2±11.2 | 0.04±0.06 | 0.09±0.08 | 0.09±0.08 |
| 50 | -0.69±0.15 | 0.68±0.17 | 0.91±0.02 | 0.68±0.17 | 4.1±1.0 | 2.6±0.3 | 6.3±3.4 | 0.23±0.15 | 0.31±0.15 | 0.30±0.15 |
| 100 | -0.29±0.00 | 0.96±0.00 | 0.97±0.00 | 0.96±0.00 | 1.6±0.0 | 1.6±0.0 | 1.7±0.0 | 0.90±0.01 | **0.94±0.01** | **0.94±0.01** |
| 1000 | -0.28±0.00 | 0.96±0.00 | 0.96±0.00 | 0.96±0.00 | 1.7±0.0 | 1.6±0.0 | 1.7±0.0 | 0.87±0.01 | 0.91±0.01 | 0.91±0.01 |
| 2990 | -0.23±0.00 | 0.94±0.00 | 0.95±0.00 | 0.94±0.00 | 2.0±0.0 | 1.9±0.0 | 2.0±0.0 | 0.74±0.01 | 0.76±0.01 | 0.76±0.01 |
| Sim2 - | -0.26±0.00 | 0.92±0.00 | 0.93±0.00 | 0.95±0.00 | 2.2±0.0 | 2.2±0.0 | 1.9±0.0 | 0.64±0.01 | 0.66±0.01 | **0.82±0.00** |
| 1 | -0.35±0.00 | 0.95±0.00 | 0.96±0.00 | 0.95±0.00 | 1.8±0.0 | 1.7±0.0 | 1.9±0.0 | 0.77±0.01 | **0.82±0.00** | **0.82±0.00** |
| UKB - | -0.53 | 0.80 | 0.88 | 0.55 | 3.6 | 3.0 | 8.9 | 6.01 | 8.05 | 10.83 |
| 1 | -0.91 | 0.41 | 0.94 | 0.41 | 5.6 | 2.2 | 13.2 | 2.54 | 7.64 | 7.64 |
| 2 | -0.80 | 0.60 | 0.90 | 0.60 | 4.8 | 2.8 | 7.9 | 6.33 | 15.19 | 15.19 |
| 5 | -0.79 | 0.61 | 0.90 | 0.61 | 4.8 | 2.9 | 7.7 | 5.74 | 13.12 | 13.13 |
| 10 | -0.76 | 0.65 | 0.90 | 0.65 | 4.6 | 2.9 | 6.9 | 6.49 | 14.03 | 14.02 |
| 20 | -0.72 | 0.69 | 0.89 | 0.69 | 4.3 | 2.9 | 6.1 | 8.54 | 17.26 | 17.28 |
| 50 | -0.68 | 0.74 | 0.89 | 0.74 | 4.0 | 2.9 | 5.4 | 10.17 | **18.08** | **18.10** |
| 100 | -0.66 | 0.75 | 0.90 | 0.75 | 3.9 | 2.9 | 5.1 | 9.65 | 17.12 | 17.13 |
| 1000 | -0.59 | 0.79 | 0.89 | 0.79 | 3.7 | 2.9 | 4.5 | 7.68 | 11.62 | 11.62 |
| 2635 | -0.53 | 0.80 | 0.88 | 0.80 | 3.6 | 3.0 | 4.2 | 6.31 | 8.61 | 8.62 |

The best estimates of brain age delta are when using *δ*_2_ or *δ*_3_ with an SVD reduction to 100 components. Reducing the number of subjects to 500 gives qualitatively similar results, with the best option again being SVD reduction to 100 components (and best *δ*_2*/*3_ prediction of true delta having a correlation of 0.82).

In all cases there is strong (negative) correlation between estimated brain age delta *δ*_1_ and actual age (column 2), an undesirable feature of the most simple brain age modelling. This is at its worst for poor models (very low SVD dimensionality), which in effect is like Fig. 1A (regression dilution). We do not show these correlations for *δ*_2_ and *δ*_3_ as these are zero by design. While many of these analyses show very good correlation between actual age and predicted age (columns 3-5), it is clear (from the final 3 columns) that this is not a good indicator of successful modelling of the brain age *δ*.

When *X* is orthogonalised via SVD, *δ*_2_ and *δ*_3_ become identical (apart from scaling factor *k*), and results improve for sufficiently large values of *J*. The best results are achieved when setting the SVD dimensionality *J* to be the “correct” number (here 100). Underestimating dimensionality (e.g., here *J* =50) damages estimation much more than overestimation (e.g., *J* =1000), an important result to keep in mind when choosing *J*.

The estimates of mean absolute values of *δ* largely show the expected pattern of results: smaller *δ* in general indicates more succesful modelling (as judged by the gold standard metric, the accuracy of recovering the true *δ*). This is expected because successful modelling of age implies relatively small modelling <u>res</u>iduals, upon which delta is based. However, there are clearly exceptions to this, e.g., with the smallest (across different models) 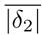 corresponding to total modelling failure (SVD *J* =1).^4^ More importantly (in practice), the optimal modelling choices can show slightly “supoptimal” mean absolute *δ* (see UK Biobank results discussed below). Hence it is important to realise that simply choosing (or tuning) a method that minimises estimated *δ* does not guarantee to give the best overall results.

While *δ*_2_ and *δ*_3_ are identical^5^ (up to a scaling factor) when using SVD, their overall scaling performs quite differently, with the scaling of *δ*_2_ being more stable and accurate overall. This means that correlations between *δ*_2_ and *δ*_3_ with true *δ* are the same, but, because of the difference in overall scaling, once these are added to true age to form estimated brain age, that will differ between the two models (as seen in r(*Y*, *Y*_*B*1_) vs. r(*Y*, *Y*_*B*2_)).

#### Simulation 2

Simulation 2 reflects a much simpler scenario (probably unrealistically simple), in order to illustrate what happens if there are no other structured effects in the data apart from age dependency. In this case the 99 structured effects were removed, and noise standard deviation raised to 10. When using the original *X* (i.e., no SVD), *δ*_2_ performs signinficantly worse than *δ*_3_. Using SVD gives results that are as good as when using *δ*_3_ with no SVD (and using *J >* 1 gives almost identical results; using *J* = 1 works well here because of the true data rank being 1). Most importantly, the simple common approach (*δ*_1_) still results in suboptimal estimation of true delta, and strong incorrect correlation with age.

#### Real data

The real data evaluation used IDPs from 19000 subjects in UK Biobank. We used 2641 IDPs spanning a range of structural, diffusion and fMRI phenotypes, to create *X*. Confounds were removed from the data as done in [Miller et al., 2016, Elliott et al., 2018] (although of course age-dependent confounds were not removed from *X*). While it is common (and generally sensible) to derive brain-aging models from healthy subjects only, and then apply those models to all subjects (including those with disease), we did not remove individual subjects from the modelling here using UK Biobank data, given that the fractions of imaged subjects having specific existent diagnoses are low (less than 10% having mental health or neurological diagnoses).

The methods described above were applied to estimate brain age delta. As this is real data we do not know the “true” delta, and therefore as a surrogate for this, for the “*Q*” columns, we instead estimated the significance of the correlation between estimated deltas and 5792 non-imaging variables, after applying the same deconfounding (though now including age-dependent confounds) to those variables. In general we would hope that “higher correlation is better”: accurately estimated brain age delta should have stronger correlations with interesting non-imaging variables such as health outcomes and biophysical markers. In order to turn the 5792 correlations into a single summary statistic, with emphasis on stronger associations rather than weakest (null) associations, we took the 99th percentile of −*log*_10_(*P*) over all non-imaging variables as our measure of success (“*Q*”).

Doing SVD reduction with *J* ≈ 50 components gave the best results, and *δ*_2*/*3_ gave much stronger associations with non-imaging variables than *δ*_1_. Encouragingly, a relatively wide range of *J* (e.g., 20-100) all gave similar strong results. The “symmetry” of *Q* (as a function of *J*, either side of the optimal dimensionality) for real data is different from the pattern found in the simulated data, where success fell off very quickly as dimensionality was reduced to be lower than the true dimensionality. We hypothesised that this was because the simulated data had a true strong cutoff in the eigenspectrum, whereas real data has a much more gradually falling eigenspectrum, as structured signals are of varying strength. We re-ran Simulation 1, this time with the strengths of the added 99 dimensions of structured subject covariation being modulated by a random number uniformly drawn from 0:1. The results were unchanged with respect to the optimal dimensionality (still being 100, and having *Q*=0.97). However, now the results were more symmetric about this, i.e., lower dimensionalities did not perform as badly, with even a low *J* of 25 giving *Q*=0.8.

The real data results are illustrated further in Fig. 3, showing clearly the value in correcting the estimate of brain age delta, and the danger of computing associations between biased brain age delta and non-imaging variables that have not been deconfounded for age.

**Figure 3:**
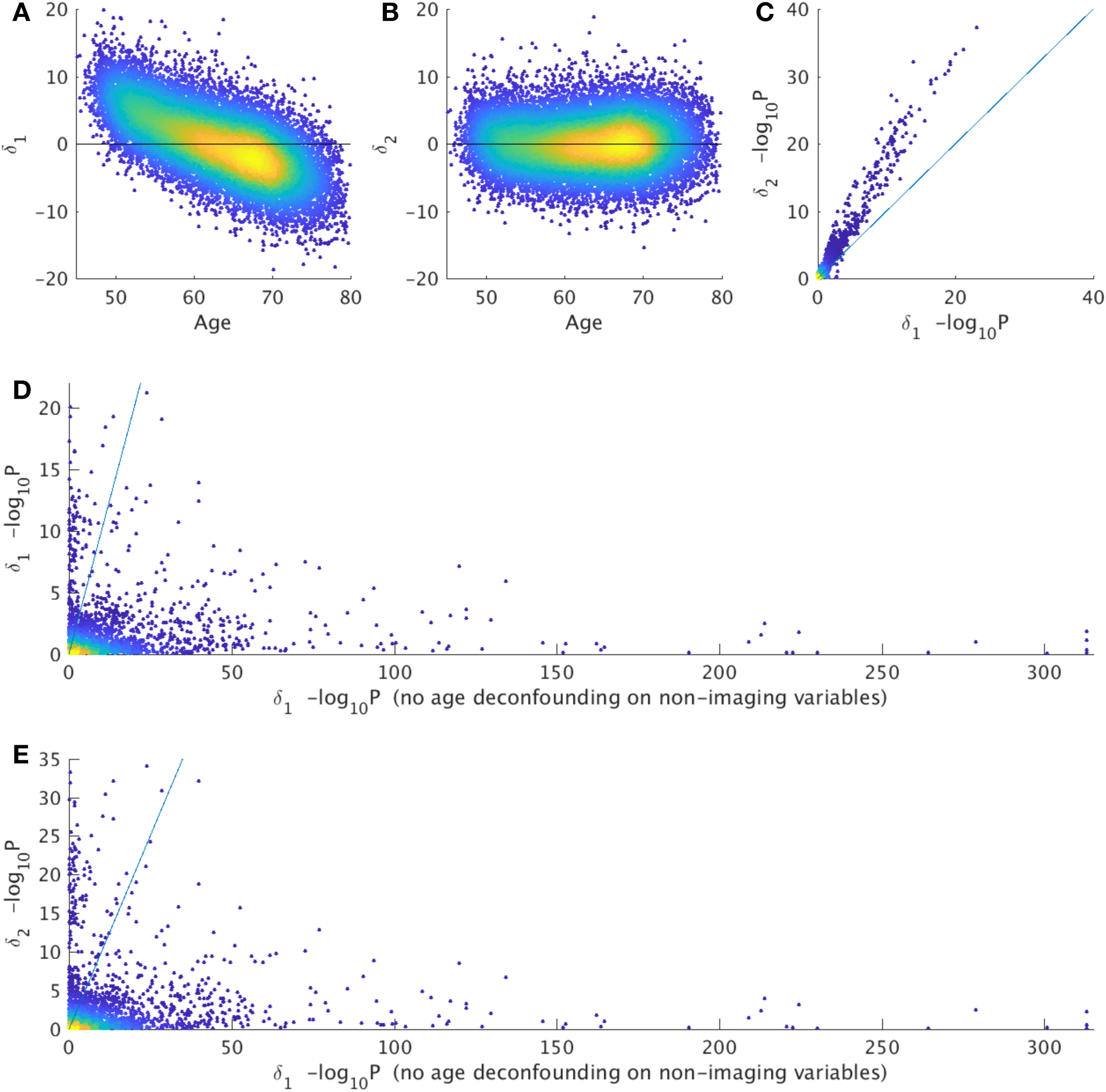
Brain age prediction results with real data from UK Biobank (N=19000). **A,B)** The first two scatterplots show the clear bias (age-dependence) in *δ*_1_ being corrected in *δ*_2_ (each point is one subject). **C)** shows relative strengths of correlations of *δ*_1_ and *δ*_2_ with (fully deconfounded) non-imaging variables, with the latter showing consistently stronger associations (each point relates to the association of a single non-imaging variable to brain age delta). P-values are uncorrected for multiple comparisons. **D)** shows the change in *δ*_1_, without (x axis) vs. with (y axis) age-deconfounding of the non-imaging variables; this illustrates the danger of looking for associations between biased brain age delta and non-imaging variables that have not been corrected for age dependence. **F)** shows a similar story, when using *δ*_2_ on the y axis; bias can cause (probably unwanted and misleading) associations (points under the y=x line), while applying the corrections described here can raise sensitivity to finding (what are hopefully valid) associations in other cases (points above the line).

### 2.7 Nonlinear relationships between age and imaging data

One cannot assume that the effect of aging on imaging measures is a linear function of age; indeed, acceleration of the effects of aging (in older age) seems quite likely, particularly in disease. The above models are simple to adapt to include an additive nonlinear term in *Y*, the most natural extension being to add a quadratic term.^6^

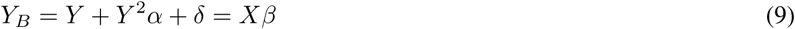

To adapt the first model described above, we can integrate the quadratic correction into the second step (that also corrects for bias in estimating the linear effect in step 1). Hence we subsume the quadratic term into the initially estimated *δ* (which is particularly straightforward if we assume that the quadratic term has been orthogonalised with respect to *Y*):

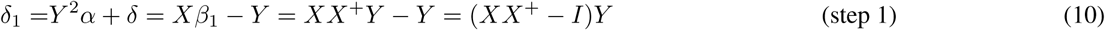

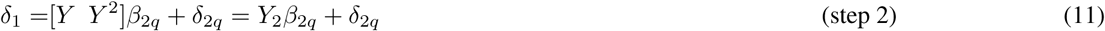

where *β*_2*q*_ has two free parameters, covering the linear and quadratic regressors. Note that the first step is identical to what we had before; hence the unchanged subscript in *δ*_1_. This can all be combined:

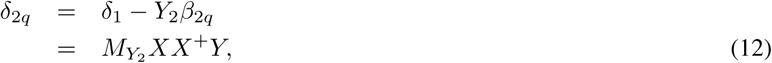

where 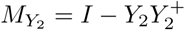.

Fig. 4 uses a simple simulation to illustrate the two stages of the model fitting described above. In **A** and **B** the biases in the one-stage modelling are apparent, and these are removed in **D** and **E**. Furthermore, the correlation of estimated delta with the true delta is improved in the second step.

**Figure 4:**
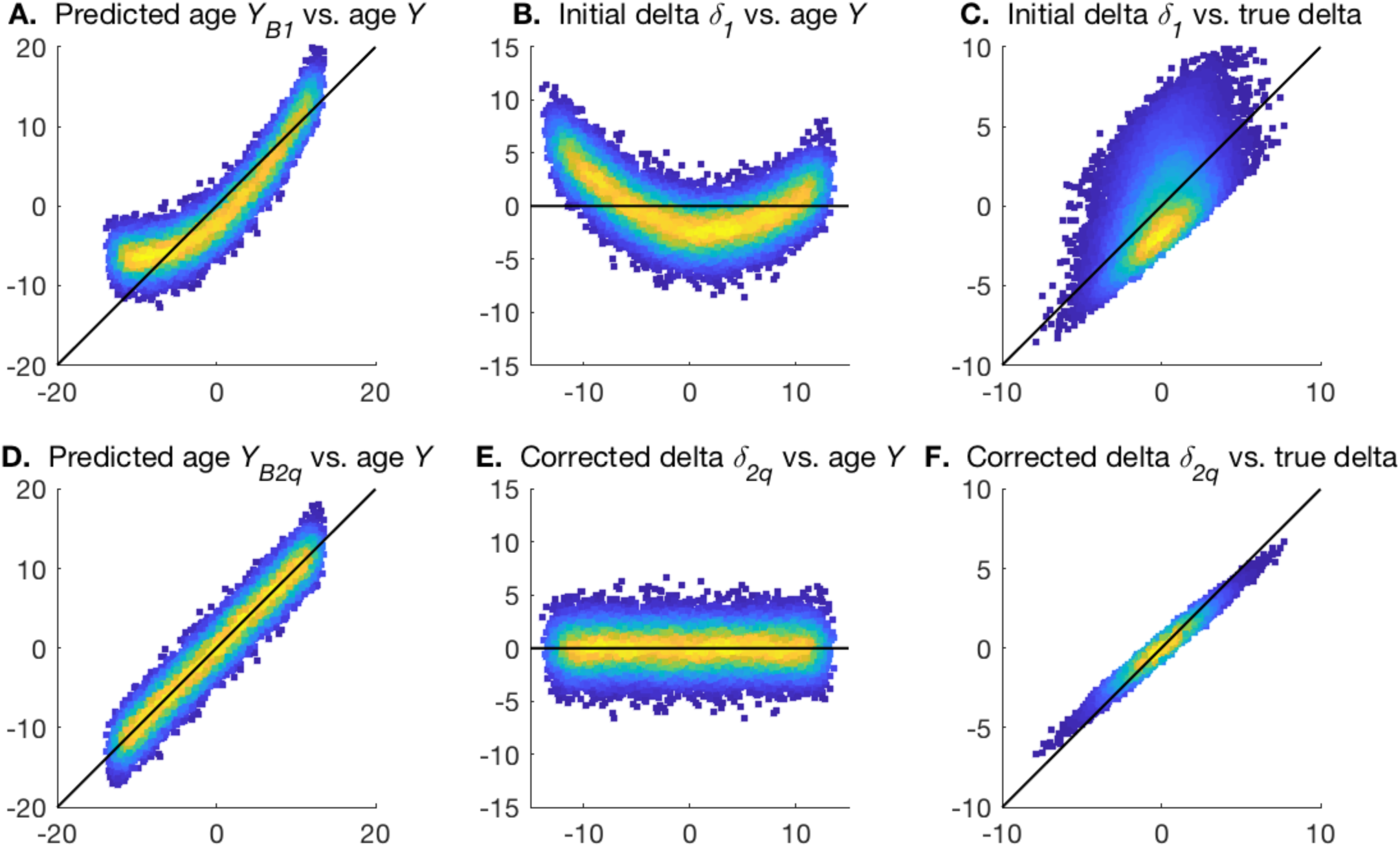
Examples of the two stages of quadratic-fit age prediction. The black lines show the ideal linear model fits. **A,D**. The plots of predicted brain age from steps 1 and 2, vs. actual age. **B,E**. The plots of estimated brain age delta from steps 1 and 2, vs. actual age. **C,F**. The plots of estimated brain age delta from steps 1 and 2, vs. true delta.

Note that step 1 above assumes that it is most useful to group the quadratic term into the residuals of the first model fitting. It may be that some study populations have a linear age effect that is smaller than the quadratic, for example with young healthy adults around the peak of the lifespan “growth-aging inverted-U curve”. In such cases, it might be possible that step 1 should fit to the quadratic term rather than the linear, with step 2 being unchanged.

For the second model, that switches predictor matrix and age, again the extension to a quadratic term in *Y* is straightforward:

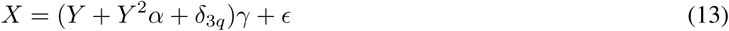

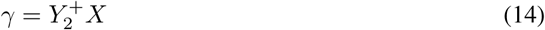

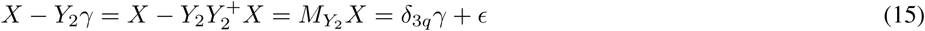

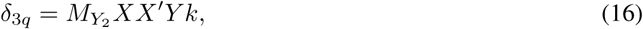

where *γ* becomes two rows of parameters (and hence incorporating *α*), and for the final line we have post-multiplied by just the first row of *γ*′, i.e., *X*^*t*^*Y k*. According to the model in Eq. 13, the two rows of *γ* should be identical (up to a scale factor), but in practice there will normally be much less variance in the data associated with the quadratic term, and we find indeed that results are improved by only using the linear term (i.e., using the first row of *γ*). (Again, in some populations, it may be better to use the quadratic part of the model rather than the linear, for the estimate of *γ*.)

Table 2 shows quantitative results from simulations of nonlinear brain aging, and from the UK Biobank real data. For the simulations, we start with the first linear simulation above, and add quadratic aging effects. For these 3 simulations we set true quadratic-term *α* values (Eq. 9) of 0, 0.01 and 0.025 respectively, corresponding to total deviations away from linear brain aging of 0, 4y and 10y.

**Table 2:** Results from quantitative evaluations of brain age estimation in the presence of nonlinear brain aging. We show results for 3 realistic simulations and a real dataset. The 3 simulations have different strengths of nonlinearity in brain age *Y* ^2^*α*, with *α* = 0, 0.01, 0.025 respectively. Different rows are for different SVD dimensionality reductions, with the first in each experiment being with no use of SVD. We show mean absolute value of different brain age *δ* estimates (for the true *δ* in the simulations, it is 1.6y) and then correlations between estimated and true *δ* (for the simulations) and significance of associations with non-imaging variables (for real data), as above.

| SVD $J$ | | $\vert \delta_1\vert $ | $\vert \delta_2\vert $ | $\vert \delta_3\vert $ | $\vert \delta_{2q}\vert $ | $\vert \delta_{3q}\vert $ | $Q(\delta_1)$ | $Q(\delta_2)$ | $Q(\delta_3)$ | $Q(\delta_{2q})$ | $Q(\delta_{3q})$ |
| --- | --- | --- | --- | --- | --- | --- | --- | --- | --- | --- | --- |
| Sim1 | - | 2.0±0.0 | 1.9±0.0 | 2.0±0.1 | 1.9±0.0 | 2.0±0.1 | 0.74±0.01 | 0.76±0.01 | 0.80±0.03 | 0.76±0.01 | 0.80±0.03 |
|  | 10 | 5.7±0.7 | 1.6±0.6 | 20.1±8.3 | 1.6±0.6 | 20.1±8.3 | 0.04±0.07 | 0.10±0.09 | 0.10±0.09 | 0.10±0.09 | 0.10±0.09 |
|  | 50 | 4.0±0.9 | 2.6±0.3 | 5.9±3.0 | 2.7±0.3 | 6.0±2.9 | 0.24±0.15 | 0.31±0.15 | 0.31±0.15 | 0.31±0.15 | 0.31±0.15 |
|  | 100 | 1.6±0.0 | 1.6±0.0 | 1.7±0.0 | 1.6±0.0 | 1.7±0.0 | 0.90±0.01 | **0.94±0.01** | **0.94±0.01** | **0.94±0.01** | **0.94±0.01** |
|  | 1000 | 1.7±0.0 | 1.6±0.0 | 1.7±0.0 | 1.6±0.0 | 1.7±0.0 | 0.87±0.01 | 0.91±0.01 | 0.91±0.01 | 0.91±0.01 | 0.90±0.01 |
|  | 2990 | 2.0±0.0 | 1.9±0.0 | 2.0±0.0 | 1.9±0.0 | 2.0±0.0 | 0.74±0.01 | 0.76±0.01 | 0.76±0.01 | 0.76±0.01 | 0.76±0.01 |
| Sim2 | - | 2.0±0.0 | 2.0±0.0 | 2.0±0.1 | 2.0±0.0 | 2.0±0.1 | 0.72±0.01 | 0.74±0.01 | 0.78±0.03 | 0.75±0.01 | 0.79±0.03 |
|  | 10 | 5.9±0.5 | 1.4±0.7 | 25.4±13.1 | 1.4±0.7 | 25.4±13.1 | 0.03±0.04 | 0.08±0.06 | 0.08±0.06 | 0.08±0.06 | 0.08±0.06 |
|  | 50 | 4.3±1.0 | 2.6±0.3 | 7.0±3.7 | 2.6±0.3 | 7.0±3.7 | 0.20±0.14 | 0.27±0.14 | 0.27±0.14 | 0.27±0.14 | 0.27±0.14 |
|  | 100 | 1.7±0.0 | 1.6±0.0 | 1.7±0.0 | 1.6±0.0 | 1.7±0.0 | 0.87±0.01 | 0.92±0.01 | 0.92±0.01 | **0.94±0.01** | **0.93±0.01** |
|  | 1000 | 1.7±0.0 | 1.7±0.0 | 1.8±0.0 | 1.6±0.0 | 1.8±0.0 | 0.85±0.01 | 0.89±0.01 | 0.89±0.01 | 0.91±0.01 | 0.90±0.01 |
|  | 2990 | 2.0±0.0 | 2.0±0.0 | 2.1±0.0 | 2.0±0.0 | 2.1±0.0 | 0.72±0.01 | 0.74±0.01 | 0.74±0.01 | 0.75±0.01 | 0.75±0.01 |
| Sim3 | - | 2.2±0.0 | 2.2±0.0 | 2.2±0.1 | 2.0±0.0 | 2.0±0.1 | 0.64±0.01 | 0.66±0.01 | 0.72±0.02 | 0.71±0.01 | 0.79±0.02 |
|  | 10 | 5.9±0.3 | 1.4±0.6 | 24.7±11.7 | 1.4±0.6 | 24.7±11.7 | 0.03±0.03 | 0.08±0.04 | 0.07±0.05 | 0.07±0.05 | 0.07±0.05 |
|  | 50 | 4.2±0.9 | 2.7±0.2 | 6.3±2.8 | 2.6±0.2 | 6.2±2.8 | 0.20±0.10 | 0.28±0.10 | 0.28±0.10 | 0.28±0.11 | 0.28±0.11 |
|  | 100 | 1.8±0.0 | 1.7±0.0 | 1.9±0.0 | 1.5±0.0 | 1.7±0.0 | 0.78±0.01 | 0.82±0.00 | 0.82±0.00 | **0.94±0.00** | **0.93±0.00** |
|  | 1000 | 1.9±0.0 | 1.8±0.0 | 2.0±0.0 | 1.6±0.0 | 1.8±0.0 | 0.75±0.01 | 0.79±0.00 | 0.79±0.00 | 0.89±0.00 | 0.89±0.00 |
|  | 2990 | 2.2±0.0 | 2.2±0.0 | 2.3±0.0 | 2.0±0.0 | 2.2±0.0 | 0.64±0.01 | 0.66±0.01 | 0.66±0.01 | 0.71±0.01 | 0.71±0.01 |
| UKB | - | 3.6 | 3.0 | 8.9 | 3.0 | 8.9 | 6.01 | 8.05 | 10.83 | 8.05 | 10.95 |
|  | 1 | 5.6 | 2.2 | 13.2 | 2.2 | 13.2 | 2.54 | 7.64 | 7.64 | 7.71 | 7.78 |
|  | 2 | 4.8 | 2.8 | 7.9 | 2.8 | 7.9 | 6.33 | 15.19 | 15.19 | 15.26 | 15.29 |
|  | 5 | 4.8 | 2.9 | 7.7 | 2.8 | 7.6 | 5.74 | 13.12 | 13.13 | 13.17 | 13.26 |
|  | 10 | 4.6 | 2.9 | 6.9 | 2.9 | 6.9 | 6.49 | 14.03 | 14.02 | 14.08 | 14.11 |
|  | 20 | 4.3 | 2.9 | 6.1 | 2.9 | 6.1 | 8.54 | 17.26 | 17.28 | 17.26 | 17.32 |
|  | 50 | 4.0 | 2.9 | 5.4 | 2.9 | 5.4 | 10.20 | 18.19 | 18.19 | **18.20** | **18.23** |
|  | 100 | 3.9 | 2.9 | 5.1 | 2.9 | 5.1 | 9.65 | 17.12 | 17.13 | 17.20 | 17.00 |
|  | 1000 | 3.7 | 2.9 | 4.5 | 2.9 | 4.5 | 7.68 | 11.62 | 11.62 | 11.61 | 11.63 |
|  | 2635 | 3.6 | 3.0 | 4.2 | 3.0 | 4.2 | 6.31 | 8.61 | 8.62 | 8.62 | 8.49 |

With no simulated nonlinear effect (Sim1), the quadratic models give identical results to the linear models (i.e., there is no noticeable penalty being paid for the additional model flexibility, presumably because, given the simplicity of the corrections to delta, overfitting is negligible). When a quadratic effect is included in the simulation, the quadratic-correction models (i.e., generating *δ*_2*q*_ and *δ*_3*q*_) provide a major improvement in accuracy over the linear models. Accuracy of recovering the true *δ* remains high even for large amounts of nonlinear behaviour, providing *J* is optimal. As before, optimal SVD data reduction outperforms using the original data matrix *X*.

The quadratic modelling in the UKB real data accounts approximately for a 2y total deviation from the linear fit. Including this quadratic modelling results in improvements in the strength of associations with non-imaging variables, although these improvements are here very small. In disease populations this might be expected to be much greater. Where the nonlinear effect is greater, it is straightforward (particularly for the first, two-stage, model) to extend the above formulations to higher powers, or, possibly more well-conditioned, nonlinear modelling such as with splines.

In all cases the “common” approach (*δ*_1_) performs significantly worse than these models that correct for bias and nonlinearity in brain age prediction.

### 2.8 Non-additive brain age delta

A different model for brain aging would be for each person to be aging at a different rate, meaning that their delta (according to the above models) is changing with age, as opposed to being fixed. Unfortunately, this cannot trivially be distinguished from having constant delta, at the level of the individual subject, if there is only one measurement (timepoint) per subject.

This ambiguity arises because subject-specific rate of aging would be considered to create a spread of trajectories around the population mean aging curve. By this definition, the effect of interest is orthogonal to the population mean aging effect, and therefore gets included in estimates of delta given above. Hence, the above aging models do not explicitly identify multiplicative brain aging, yet readily fit data from individual subjects at a single timepoint as an additive offset term regardless of the cause (constant offset vs. multiplicative).

However, while additive and multiplicative brain aging cannot easily be disambiguated at the subject level, it is possible to apply the above models, and then use the resulting delta values to estimate how scaling of the size of delta is changing with age in the population as a whole:

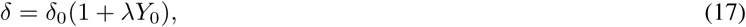

where we form a temporary version of age *Y*_0_, which is a linear mapping of *Y* into the range 0:1, and hence |*δ*_0_|relates to the brain age delta distribution for the youngest subjects in the data. The product on the right is a pointwise product between the two column vectors *δ*_0_ and (1 + *λY*_0_). The reason for formulating things this way will now become clear, where we solve by squaring, taking the natural log, approximating the log function on the right using a power series expansion, and solving with linear regression:

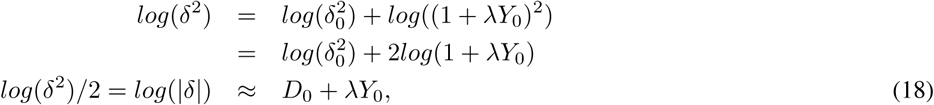

where 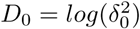 is the noise in this model and this assumues that *λ* is not very large and negative. Hence fitting the model *Y*_0_ to data *log*(|*δ*|) gives us an initial *λ*, which we can then adjust for the expansion approximation error:

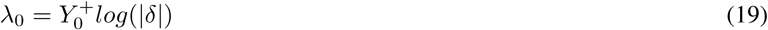

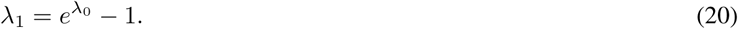

To evaluate this with simulated data, we took the first linear simulation described above, and added age-dependence by setting true lambda to 0.5 according to Eq. 17. We then ran the simulation 10 times, with *J* =100, estimating *λ* from *δ*_2_. This resulted in estimated *λ* = 0.46 ± 0.09.

On real data from UK Biobank, using 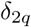 estimated using optimal SVD setting of *J* = 50, we find that *λ*=0.13, with the regression-based fitting also estimating this as being significantly greater than zero (*P* =0.001). This corresponds to an increase in “spread” of brain age delta of about 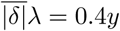 when moving from the youngest to oldest subjects in UKB (45 to 80y). This provides evidence for a modest non-additive brain aging effect in this largely healthy, aging population.

### 2.9 Further results from UK Biobank brain age estimation

We carried out brain age delta estimation on UK Biobank data as described above, and also for females and males separately. Below we report results using 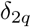.

The delta estimations for females were almost identical when comparing delta estimated purely for females vs. estimated from the all-in-one delta estimation from females and males together (*r*=0.98). The same was true for males (*r*=0.97). From the all-in-one analysis, females had a mean brain age delta that was 0.7y higher than in males.

We then correlated these various versions of brain age delta against 5792 non-imaging variables from UK Biobank, and converted these uncorrected P-values into −*log*_10_*P*. As expected from the above results, these vectors of correlation significance (with one entry in the vector for each non-imaging variable) were highly similar when comparing sex-separated delta estimation for females against when taking the deltas for females from the all-in-one estimation (*r*=0.98). The same was found for males (*r*=0.98). Hence, for sex-specific results reported below, we only list those from the sex-separated delta estimations.

Comparing −*log*_10_*P* derived from all subjects (both sexes) from the all-in-one analysis against the sex-separated estimates of −*log*_10_*P* showed greater differences (all subjects vs. female: *r*=0.87, vs. male: *r*=0.73). Comparing −*log*_10_*P* derived from female-only analysis against male-only gave *r*=0.36.

The strongest associations with non-imaging variables are shown in Tables 3 and 4.^7^ Bonferroni correction, for the number of non-imaging variables, gives a threshold for −*log*_10_*P* of 5.1, but to limit the number of reported associations to a reasonable degree (and because effect sizes become tiny at this threshold, the weakest passing this being 0.1% variance explained!), we report results for −*log*_10_*P >* 8. The non-imaging variables are denoted via unique variable codes and brief descriptions. For more information on any variable, one can take the initial integer from the ID, and search for the variable on the UK Biobank website: https://biobank.ctsu.ox.ac.uk/showcase/search.cgi

**Table 3:** The strongest associations between brain age delta and non-imaging variables in UK Biobank, 19038 subjects, males and females combined. Positive correlations (*red italics*) imply accelerated brain aging.

| $-\log_{10}P$ | r | variable ID | variable description |
| --- | --- | --- | --- |
| 37.5 | -0.10 | 4125-2.0 | Heel bone mineral density |
| 34.5 | 0.09 | 4080-0.1 | *Systolic blood pressure* |
| 32.0 | -0.09 | 21002-2.0 | Weight |
| 32.0 | 0.09 | 4079-0.1 | *Diastolic blood pressure* |
| 29.1 | -0.18 | 23236-2.0 | Total BMD (bone mineral density) |
| 26.7 | -0.08 | 21001-2.0 | Body mass index (BMI) |
| 26.4 | -0.08 | 49-2.0 | Hip circumference |
| 25.1 | 0.08 | 20116-0.0 | *Still smoking* |
| 25.0 | 0.08 | 30040-0.0 | *Mean corpuscular volume* |
| 24.0 | 0.08 | 12702-2.0 | *Cardiac index during PWA* |
| 23.3 | 0.08 | 12687-2.1 | *Mean arterial pressure during PWA* |
| 23.2 | -0.16 | 23235-2.0 | Total BMC (bone mineral content) |
| 21.9 | -0.07 | 23120-0.0 | Arm fat mass (right) |
| 21.9 | 0.07 | 30050-0.0 | *Mean corpuscular haemoglobin* |
| 20.5 | 0.08 | 12673-2.0 | *Heart rate during PWA* |
| 18.6 | 0.07 | 12682-2.0 | *Cardiac output during PWA* |
| 18.6 | 0.07 | 102-2.0 | *Pulse rate* |
| 18.6 | -0.14 | 23226-2.0 | Head BMD (bone mineral density) |
| 18.0 | 0.07 | 1558-2.0 | *Alcohol intake frequency* |
| 16.2 | -0.06 | 20015-2.0 | Sitting height |
| 16.0 | -0.06 | 30010-0.0 | Red blood cell (erythrocyte) count |
| 15.8 | -0.06 | 23105-0.0 | Basal metabolic rate |
| 15.0 | 0.06 | 137-2.0 | *Number of treatments/medications taken* |
| 14.3 | 0.06 | 2443-2.0 | *Diabetes diagnosed by doctor* |
| 14.0 | 0.06 | 1249-2.0 | *Past tobacco smoking* |
| 13.8 | 0.06 | 30270-0.0 | *Mean sphered cell volume* |
| 13.0 | 0.08 | 20157-0.0 | *Duration to complete alphanumeric path (cognition)* |
| 12.3 | -0.05 | 23099-0.0 | Body fat percentage |
| 11.4 | -0.05 | 20016-2.0 | Fluid intelligence score (cognition) |
| 10.7 | 0.05 | 404-2.7 | *Duration to first press of snap-button (cognition)* |
| 10.2 | -0.06 | 20195-0.0 | Number of symbol digit matches attempted (cognition) |
| 9.9 | -0.06 | 20159-0.0 | Number of symbol digit matches made correctly (cognition) |
| 9.9 | -0.05 | 3064-2.1 | Peak expiratory flow (PEF) |
| 9.8 | -0.09 | 22408-2.0 | Abdominal subcutaneous adipose tissue volume |
| 9.5 | -0.05 | 1458-2.0 | Cereal intake |
| 9.4 | 0.05 | 20023-2.0 | *Mean time to correctly identify matches (cognition)* |
| 9.3 | 0.05 | 1970-0.0 | *Nervous feelings* |
| 8.7 | -0.05 | 738-2.0 | Average total household income before tax |
| 8.7 | 0.06 | 20133-0.1 | *Time to complete round, pairs matching (cognition)* |
| 8.6 | -0.04 | 709-0.0 | Number in household |
| 8.4 | 0.05 | 1588-0.0 | *Average weekly beer plus cider intake* |
| 8.2 | 0.05 | 1568-0.0 | *Average weekly red wine intake* |

**Table 4:**
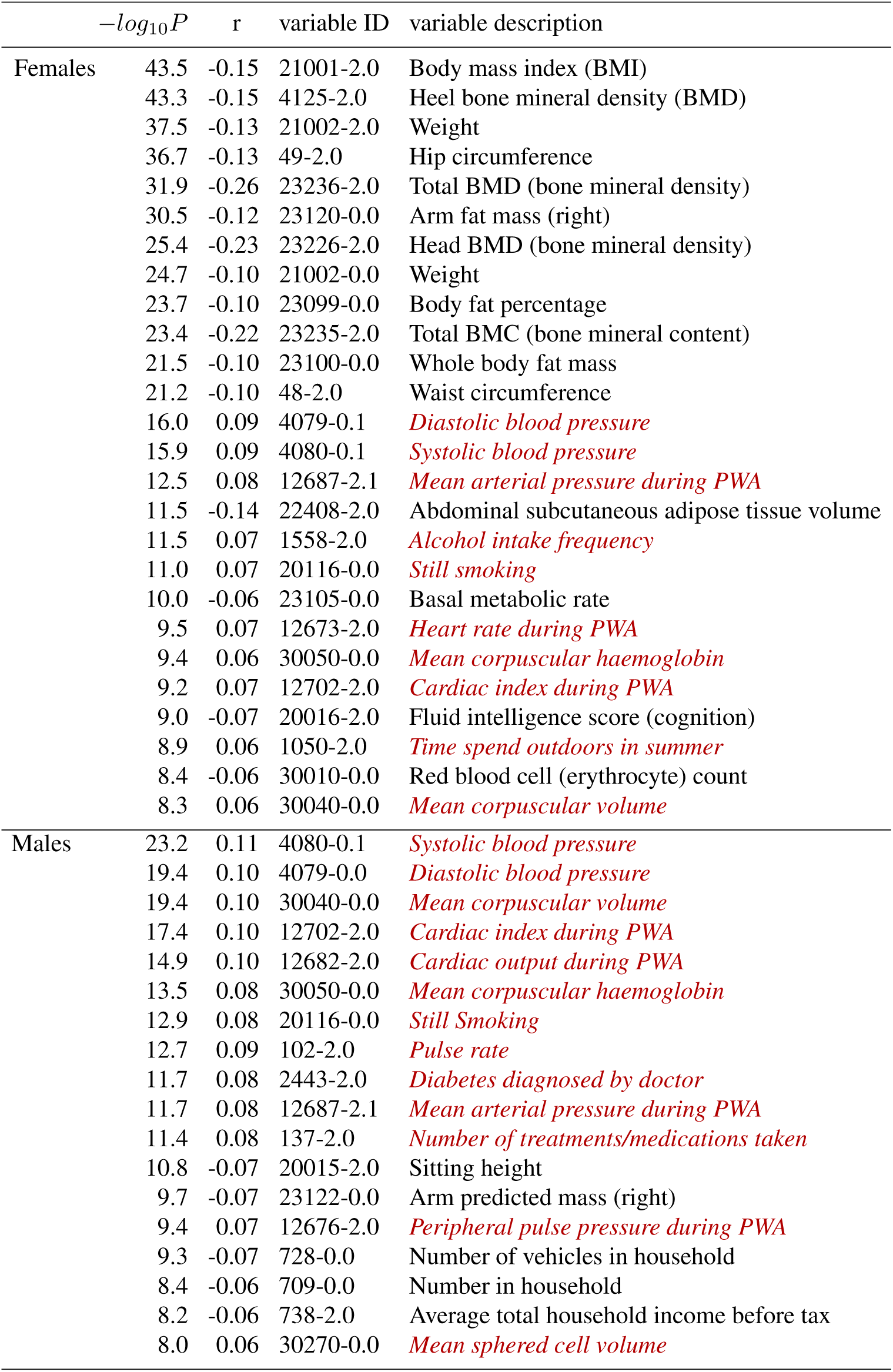
The strongest associations between brain age delta and non-imaging variables in UK Biobank, for just the 10112 females, and just the 8926 males. Positive correlations (*red italics*) imply accelerated brain aging.

For these lists we have manually removed largely-redundant (highly similar) variables for purposes of readability, listing just the strongest associated result in each case. If a variable is positively correlated with brain age delta, this implies that accelerated brain aging is associated with larger values of that variable, i.e., it is a “bad” life factor or biological measure. Of course, the results do not allow one to infer causality.

There are several strong patterns that emerge. Higher body weight, body fat and bone density are all associated with reduced brain aging, largely in females. Higher blood pressure, heart rate and blood haemoglobin are all associated with accelerated brain aging, in both females and males, but to a slightly stronger extent in males.

Smoking and alcohol are associated with accelerated brain aging. There are several life-factor/life-style measures associated with delta, presumably reflecting socio-economic status, where the biological causality is likely complex (e.g., household income, number of vehicles in household, and possibly also related, time spent outdoors in summer). Several measures from the cognitive testing are associated with brain age delta, in all cases in the direction one might expect; measures of success/accuracy correlate negatively, and measures of “time taken” (to complete a cognitive task) correlate positively.

Finally, two associations related to clinical diagnosis/treatment, both found in males, are that the number of treatments/medication taken, and diagnoses of diabetes, are both associated with accelerated brain aging.

We also looked at the correlations between delta and the imaging features (IDPs). This is expected to largely reflect which IDPs most strongly contribute to the modelling of the brain age delta, but of course, being a univariate analysis is straightforward to interpret and does not take into account redundancy across IDPs. Table 5 lists these, sorted according to decreasing strength of correlations computed from all subjects (females and males), but also showing correlations for just females and just males. We do not report *P* here, as, given the relatively strong correlations, the *P* values are too strongly significant to be differentially informative. IDPs are included in the table if any of the 3 correlations (from females only, males only, or all subjects) are stronger than 0.3.

**Table 5:**
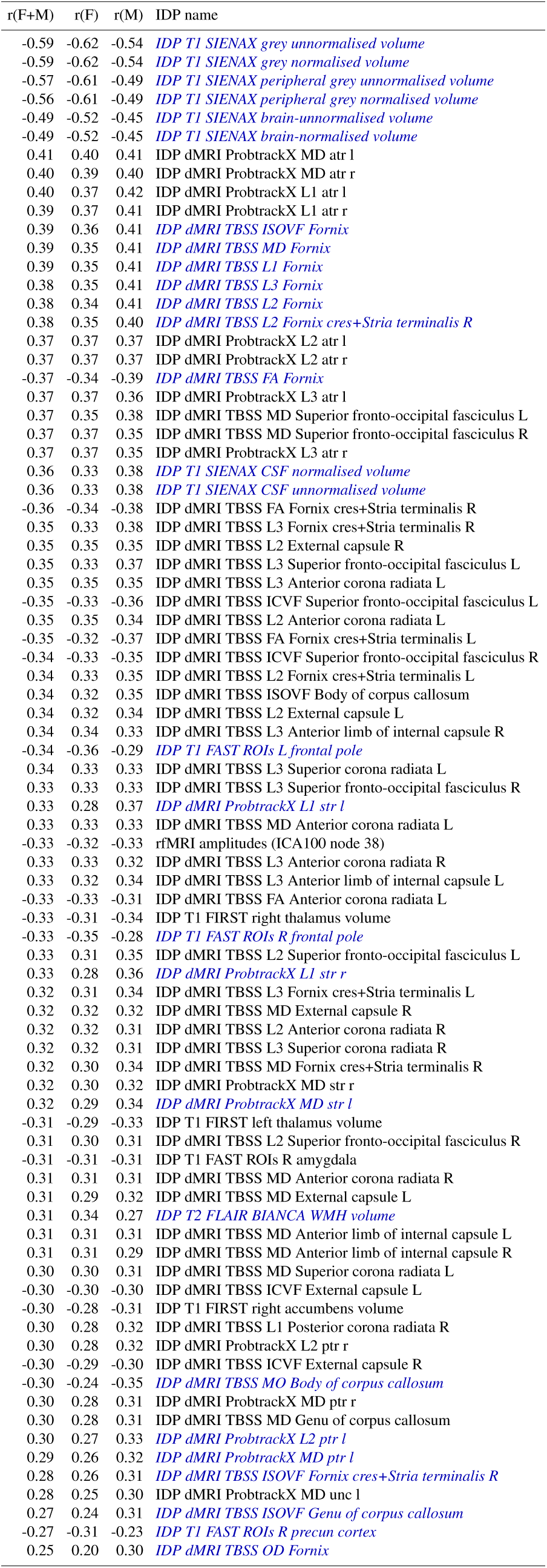
The strongest associations between brain age delta and imaging-derived phenotypes (IDPs) in UK Biobank, for all subjects, and for just the 10112 females, and just the 8926 males. IDP names listed in *blue italics* denotes where the correlations differ between females and males by more than 0.05.

We highlight in blue the IDPs for which the correlation with brain age delta is different between females and males by more than 0.05. More detailed descriptions of the IDPs (including expansions of some of the anatomical acronyms) can be found here; http://www.fmrib.ox.ac.uk/ukbiobank/IDPinfo_Jan2018.txt

The strongest associations with delta are for grey and white total volumes, both normalised for head size and unnormalised. (Note also that we have used fully deconfounded versions of the IDPs for these correlations with delta, which included regressing out head size). There are then many measures of white matter microstructure (derived from the diffusion MRI) that correlate with delta, a minority of which are reasonably strongly different between the sexes.

## 3 Conclusions

We have discussed various problems and potential solutions relating to the estimation of brain age delta. It has commonly been the case that estimated delta is not orthogonal to age, and this can result in false associations with non-imaging variables, as well as a loss in sensitivity to finding valid associations. While estimation bias is a common general statistical issue, it is specifically problematic in the area of brain-age modelling, because the final computed delta is the difference of the estimated quantity and the true value.

We found that the strength of correlation of estimated brain age with actual age is not guaranteed to be a good indicator of optimal delta estimtion, and neither is the (smallness of) the size of mean absolute delta.

We have shown that brain age delta can be estimated well by fitting one of the models described in this paper (e.g., see algorithm below), where bias in brain age estimation, and nonlinear dependence can be adjusted for.

We have also described how one might study non-additive effects of aging, where different subjects age at different rates. One of the few studies that has carried out linear age adjustment of delta is [Cole et al., 2017]. Interestingly, in this case, the adjustment reduced the strength of assocations with external information (in that case, heritability). While this goes against most of our findings above, it may relate to non-additive aging, in which case, combining the modelling (for non-additive aging) presented here with the corrected delta models presented above might optimise a delta estimation.

One simple, effective and computationally cheap tool for modelling brain age (before delta estimation) is to reduce the input data features (from the brain imaging) via SVD. We have shown that there is likely to be an optimal SVD dimensionality for the data reduction, and that it is safer to slightly overestimate rather than underestimate this dimensionality.

To apply the recommended approach for estimating brain age delta, apply the following.

1. Your vector of ages is *Y* (subjects × 1).
2. Your matrix of brain imaging measures is *X* (subjects × features/voxels).
3. Subtract the means from *Y* and all colums in *X*.
4. Use SVD to replace *X* with its top 10-25% vertical eigenvectors.
5. Compute *Y* ^2^, demean it and orthogonalise it with respect to *Y* to give 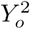.
6. Create matrix 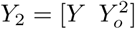.
7. The initial model is *Y*_*B*1_ = *Xβ*_1_ + *δ*_1_. Do:
  a. Compute initial age prediction *β*_1_ = *X*+*Y* giving *Y*_*B*1_ = *Xβ*_1_ (where *X*^+^ is the pseudo-inverse of *X*).
  b. Compute initial brain age delta *δ*_1_ = *Y*_*B*1_ −*Y*.
8. The corrected model is 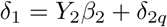 Do:
  a. Compute corrected model fit 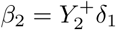 (correcting for bias in the initial fit and quadratic brain aging).
  b. Compute final brain age delta 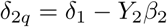

All associations of brain-age delta with UK Biobank non-imaging variables are listed in a supplementary spreadsheet. Example Matlab code for the delta computations and all simulations can be found at: http://www.fmrib.ox.ac.uk/BrainAgeDelta

## 4 Acknowledgments

We are grateful for funding from the Wellcome Trust. This research has been conducted in part using the UK Biobank Resource under Application Number 8107. We are grateful to UK Biobank for making the data available, and to all UK Biobank study participants, who generously donated their time to make this resource possible. Thanks to Saad Jbabdi, Matteo Bastiani, Elliot Tucker-Drob and Simon Cox for discussions on this work.

## Footnotes

1 We assume throughout that *Y* and all columns in *X* are demeaned (shifted to have zero mean), to simplify equations with no loss of generality. Also, for readability, we do not in general differentiate in our notation between true parameters and estimates of those parameters.

2 Alternatively, if this is poorly conditioned, e.g., with the number of features *D* too large, using penalised regression.

3 Here we are using pseudo-inverse notation even though *Y* is just a column vector.

4 In the case where *X* is a useless model, *δ*_2_ tends to zero (because *X* explains none of *Y*), whereas *δ*_3_ tends to infinity (because *Y* explains none of *X*: the fit becomes horizontal and the horizontal errors large).

5 In fact, as can been seen in the “*Q*” columns, in practice here they are not exactly identical, simply because these simulated datasets are fit in a cross-validated way, with randomly assigned fold memberships causing small variations in outcomes.

6 Without loss of generality, and to simplify notation and interpretation, in our equations and in the simulations, by *Y* ^2^ we mean a quadratic term that has been orthogonalised with respect to *Y* and demeaned. If *Y* has already been demeaned, then *Y* ^2^ will already be close to being orthogonal to *Y*.

7 Note that there is not a monotonic relationship between *r* and *P*, because of different missing-data patterns in different variables resulting in varying *N*. Also, unlike with the results correlating delta against the imaging IDPs below, here we sort on *P* rather than *r*, given that the *r* values are rather small, leading to increased relevance of *P*.

